# Variation in root secondary metabolites is shaped by past climatic conditions

**DOI:** 10.1101/2020.01.25.919654

**Authors:** Zoe Bont, Tobias Züst, Meret Huber, Matthias Erb

**Affiliations:** Institute of Plant Sciences, University of Bern, Altenbergrain 21, 3013 Bern, Switzerland; Institute of Plant Biology and Biotechnology, University of Münster, Schlossplatz 7-8, 48143 Münster, Germany

**Keywords:** climatic conditions, dandelion, environmental gradient latex, herbivore defence, intraspecific variation, multifunctionality, secondary metabolites

## Abstract

1. Plants can adapt to changing environments by adjusting the production and maintenance of diverse sets of bioactive secondary metabolites. To date, the impact of past climatic conditions relative to other factors such as soil abiotic factors and herbivore pressure on the evolution of plant secondary metabolites is poorly understood, especially for plant roots.
2. We explored associations between root latex secondary metabolites in 63 *Taraxacum officinale* populations across Switzerland and past climatic conditions, soil abiotic parameters, and root herbivore pressure. To assess the contribution of environmental effects, root secondary metabolites were measured in F0 plants in nature and F2 plants under controlled greenhouse conditions.
3. Concentrations of root latex secondary metabolites were most strongly associated with past climatic conditions, while current soil abiotic factors or root herbivore pressure did not show a clear association with root latex chemistry. Results were identical for natural and controlled conditions, suggesting heritable trait variation rather than environmental plasticity as underlying factor.
4. *Synthesis.* We conclude that climatic conditions likely play a major role in the evolution of root secondary metabolites. Direct abiotic effects are likely underlying this pattern, hinting at a novel role of root latex metabolites the tolerance of abiotic stress.

## Introduction

Plants produce a tremendous variety of structurally diverse organic compounds, so-called secondary or specialized metabolites. These metabolites defend plants against herbivores and pathogens (Mithöfer & Boland, 2012) increase abiotic stress tolerance (de Costa, Yendo, Fleck, Gosmann, & Fett-Neto, 2013; Hazarika & Rajam, 2011; Qi, Yang, Yuan, Huang, & Chen, 2015), facilitate mutualisms (Peters, Frost, & Long, 1986; Schäfer et al., 2009; Stevenson, Nicolson, & Wright, 2017), promote micronutrient uptake (Hu et al., 2018; Kobayashi & Nishizawa, 2012), or act as growth and defence regulators (Francisco et al., 2016; Kim, Ciesielski, Donohoe, Chapple, & Li, 2014; Malinovsky et al., 2017). Although secondary metabolites can have highly specialized functions, there is growing evidence that individual many of them serve multiple purposes (Hu et al., 2018; Katz et al., 2015; J. Li et al., 2018; Malinovsky et al., 2017; Møller, 2010). Multifunctional secondary metabolites help the plant to minimize the effort required for biosynthesis and compound maintenance and are therefore seen as a cost-effective resource allocation strategy (reviewed in Neilson, Goodger, Woodrow, & Møller, 2013).

Given the importance of secondary metabolites in adjusting a plant’s physiology to its continuously changing environment, abiotic and biotic factors are expected to exert strong selective pressure on plant secondary metabolism (Hartmann, 2007). Hence, metabolite profiles can vary substantially even within the same species. The unique set of secondary compounds produced by a plant population likely reflects the particular demands of the plant’s ecological niche and its current selective environment (Moore, Andrew, Külheim, & Foley, 2014). In *Arabidopsis thaliana* populations, for instance, geographic variation in the abundance of *A. thaliana* chemotypes is strongly associated with the geographic pattern in the relative abundance of two specialist herbivores (Züst et al., 2012). Patterns of secondary metabolite profiles across populations have also been directly or indirectly linked to geographical gradients such as latitude or elevation (Abdala-Roberts, Moreira, Rasmann, Parra-Tabla, & Mooney, 2016; Anstett, Ahern, Johnson, & Salminen, 2018; Coll Aráoz, Mercado, Grau, & Catalán, 2016; Moles et al., 2011; Woods et al., 2012). As changes in geographic locations are associated with substantial variation in both biotic and abiotic traits, integrated analyses that take different environmental parameters into account are helpful to identify the environmental factors that shape the evolution and expression of plant secondary metabolites in natural populations.

Intraspecific variation in secondary metabolites can be constitutive (Züst et al., 2012). and can be further amplified by phenotypic plasticity (Huber, Bont, et al., 2016). To partition this natural variation into constitutive, genetically fixed and inducible, plastic components, common garden experiments provide a useful tool (Anstett et al., 2018; Hahn, Agrawal, Sussman, & Maron, 2018; Stevens, Brown, Bothwell, & Bryant, 2016) as they control for the contribution of the variation induced by the local environment. In a common garden study with *Artemisia californica*, for instance, Pratt, Keefover-Ring, Liu, and Mooney (2014) showed that genetically-based variation in terpene composition and monoterpene concentration is associated with latitude of the source population and the corresponding differences in precipitation. Furthermore, comparing the production of secondary metabolites under controlled environmental conditions with their production in the plant’s natural habitat allows the assessment of environmental plasticity (Abdala-Roberts et al., 2016; Castillo et al., 2013). Thus, studies that measure secondary metabolites among populations in both their natural habitat and in common gardens can help determine to what extent a plant’s chemical phenotype is fixed or plastic in response to its environment (Hahn et al., 2018).

Many secondary metabolites are found in both roots and shoots of a plant, yet as a result of the distinct functions of these two plant parts (van Dam, 2009) composition and regulation of metabolites often differ above and below ground (Hartmann, 2007; Johnson, Erb, & Hartley, 2016). Changes in abiotic factors result in different morphological and physiological responses in roots compared to shoots, with different metabolic patterns that seem to allow complementary adjustments above and below ground through phenotypic plasticity (Gargallo-Garriga et al., 2015; Mithöfer & Boland, 2012; Rasmann & Agrawal, 2008). Above ground, climatic conditions of natural habitats have been shown to shape heritable intraspecific variation of secondary metabolites (Hahn et al., 2018; Pratt et al., 2014), indicating that past and present climatic characteristics select for specific chemotypes. Below ground, climatic conditions likely influence heritable variation of secondary metabolites as well, but experimental evidence for such effects is lacking so far. Furthermore, soil physical and chemical properties such as humus content or pH may affect plant physiology by determining the amount and composition of available nutrients (Dubuis et al., 2013), which in turn can result in differences in plant chemistry (Cunningham, Summerhayes, & Westoby, 1999; Meindl, Bain, & Ashman, 2013). In addition to abiotic factors, biotic factors such as herbivores have been identified to drive intraspecific variation of defensive secondary metabolites in the leaves (Agrawal, 2011; Schuman & Baldwin, 2016; Züst et al., 2012) and recently also in the roots (Huber, Bont, et al., 2016). Although the effects of specific environmental factors on plant secondary metabolites have been studied extensively, the simultaneous and combined effects of abiotic and biotic factors including climatic conditions, soil geochemical properties and herbivores on heritable variation in plant secondary metabolites remain poorly understood, especially below ground.

In this study, we investigated the role of abiotic and biotic factors in shaping variation in root secondary metabolites of the globally distributed common dandelion, *Taraxacum officinale* agg. (Asteraceae), as a model system. Dandelion accumulates toxic secondary metabolites primarily in latex, a milky, often sticky sap that is transported and stored in pressurized laticifers, to be released upon damage by herbivores. Laticifers allow for compartmentalised storage and deployment of toxic compounds while also preventing autotoxicity by the often highly reactive substances (Hagel, Yeung, & Facchini, 2008). Latex can be found in approximately 10 % of all flowering plant species and contains a rich variety of secondary metabolites (Agrawal & Konno, 2009; Castelblanque et al., 2017). Based on its physical and chemical properties, latex has been associated with defensive functions against herbivores and pathogens, and no other functions are currently known (Konno, 2011).

Our previous work revealed that latex of *T. officinale* contains three major classes of secondary metabolites: the sesquiterpene lactone taraxinic acid ß-D-glucopyranosyl ester (TA-G), hydroxyphenylacetate inositol esters with either two or three side chains (di-PIEs respectively tri-PIEs) and several triterpene acetates (TritAc) (Huber et al., 2015). Both TA-G and PIEs are involved in defence against herbivores and are highly variable among natural populations (Agrawal, Hastings, Fines, Bogdanowicz, & Huber, 2018; Bont et al., 2017; Huber, Bont, et al., 2016; Huber, Epping, et al., 2016), and we have demonstrated that the concentration of TA-G is shaped by selection from the major native root herbivore *Melolontha melolontha* (Huber, Bont, et al., 2016). However, both abiotic and biotic environmental factors can influence the quantity of latex exudation (Barton, 2014; Raj, Das, Pothen, & Dey, 2005; Woods et al., 2012) and the relative composition of the diverse latex secondary metabolite mixtures may be likewise affected by environmental conditions, although experimental evidence for this assumption is scarce.

Here, we investigated environmental drivers of intraspecific variation in root latex secondary metabolites of *T. officinale*. Focusing on the main secondary metabolites of *T. officinale* latex, we determined metabolite profiles of 63 populations growing in their natural habitat and of their offspring grown under controlled conditions in the greenhouse. We recorded and inferred current and historic root herbivore abundance as well as soil geochemical parameters in the natural habitats of the different populations. We furthermore retrieved climatic conditions of the different habitats over the last 20 years, and performed model selection to determine which environmental variables are most strongly associated with naturally expressed and heritable variation in root secondary metabolites.

## Materials and methods

### Study species

The common dandelion, *T. officinale,* is a latex-producing perennial herb native to Eurasia (Stewart-Wade, Neumann, Collins, & Boland, 2002). It is described as a species complex that consists of diploid outcrossing, triploid apomictic and, in rare cases, tetraploid individuals (Verduijn, Van Dijk, & Van Damme, 2004). In Switzerland, *T. officinale* most commonly colonizes low- and mid-elevation habitats, but can also be found at altitudes higher than 2000 m a.s.l. (Calame & Felber, 2000). The main peak flowering time of *T. officinale* in the northern hemisphere is from April to June, with plants growing in warmer habitats at lower altitude flowering earlier than plants growing in colder habitats. Approximately 10-12 days after flowering, seed maturation is completed and each capitulum produces several hundred seeds equipped with a parachute-like structure to facilitate wind dispersal (Honek & Martinkova, 2005). Using a simulation model for wind dispersal, Tackenberg, Poschlod, and Kahmen, (2003) calculated that more than 99.5% of released dandelion seeds land within 10 m distance, whereas 0.014% had the potential to be dispersed >1km. Pollination, on the other hand, occurs mainly within a range of a few meters (Lázaro & Totland, 2010; Takakura, Matsumoto, Nishida, & Nishida, 2011). Thus, although occasional long-distance seed dispersal events occur, gene flow between distant populations is likely restricted due to limitations in dispersal distances of pollen and seeds.

### Field sites and environmental data

From April to June 2016, we identified and characterized 63 *T. officinale* populations across Switzerland *in situ*. Populations were selected based on proximity to meteorological monitoring stations of MeteoSwiss, the Swiss Federal Office for Meteorology and Climatology, with all field sites being located within a maximal distance of 1 km from a station. Based on this criterion we evenly distributed the sampling sites across Switzerland, with altitudes of populations ranging from 200 – 1600 m a.s.l., (Fig. 1, Table S1). Each population was visited once towards the end of its flowering period. On each field site, we marked a square of 25 m^2^ in a representative area of the field and haphazardly selected 15 *T. officinale* plants within this square for latex and seed collection. Where available, information on land use intensity over the last decades was obtained from landowners. To characterize long-term climatic conditions, ten variables were selected from the MeteoSwiss database (Table 1). The chosen variables represent temperature, precipitation, light and air pressure conditions of the populations, which are important abiotic factors affecting plant physiology and production of secondary metabolites (Wallis, Huber, & Lewis, 2011; Zhou et al., 2017; Zidorn, 2010). Data was obtained for the years 1996 – 2015 and averaged over this period for each population.

**Figure 1.**
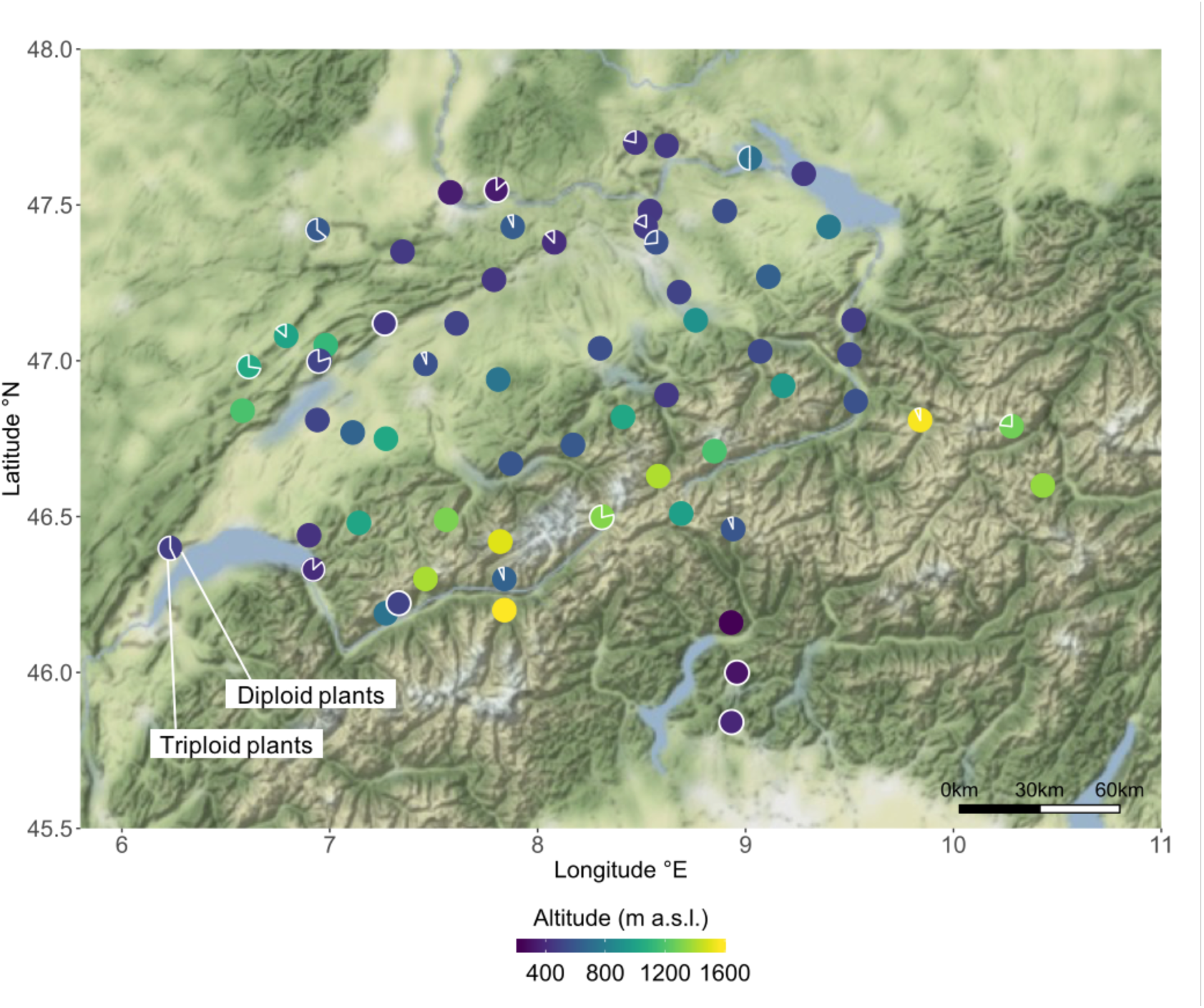
*T. officinale* sampling locations across Switzerland. Each circle displays one of the 63 sampled populations. Colour gradient of the circle fill represents the altitude of the populations. The proportion of the circle that is outlined in white colour corresponds to the percentage of triploid plants of the respective population, whereas the unmarked part of the circle corresponds to the percentage of diploids.

**Table 1.** Correlations of climatic variables with the first two principal components (climPCA1, climPCA2) from a principal component analysis of the climatic conditions of the sampled *T. officinale* populations.

| Climate variable | Description | climPCA1 | climPCA2 |
| --- | --- | --- | --- |
| AIR | Annual mean of air pressure | 0.413 | -0.113 |
| PRE | Annual precipitation | -0.026 | -0.356 |
| PRED | Days per year with precipitation | -0.145 | -0.503 |
| RAD | Annual mean of global radiation | -0.175 | 0.546 |
| SUN | Annual sunshine duration | 0.114 | 0.500 |
| TM | Annual mean temperature | 0.435 | -0.008 |
| TMN | Annual mean of the minimal temperature per day | 0.342 | -0.125 |
| TMX | Annual mean of the maximal temperature per day | 0.375 | 0.107 |
| T0D | Days per year with minimal temperature below 0°C | -0.401 | 0.112 |
| T25D | Days per year with maximal temperature over 25°C | 0.396 | 0.138 |

For soil analysis, soil samples were taken from each field site. We used a ‘Swiss Sampler’ (Eric Schweizer AG, Thun, Switzerland) to take 15 soil samples of the top 15-20 cm soil layer along one diagonal of the 25 m^2^ square. Soil samples were then pooled and stored at 4 °C after returning to the lab. Each pooled sample was then analysed for PEP (proof of ecological performance) (performed by Labor für Boden- und Umweltanalytik, Eric Schweizer AG, Thun, Switzerland). From this data we selected humus content and soil pH for further analyses because both are important determinants of the amount and composition of available nutrients for plants and thus influence plant physiology (Dubuis et al., 2013).

To assess the abundance of belowground herbivores we used two approaches: first, three of the 15 selected dandelion plants were excavated in the field, and the number of belowground herbivores in the rhizosphere and surrounding soil core (approximately 15 x 15 x 15 cm) was counted and averaged per plant. Second, we categorized the field sites belonging to either an area with historically high density of *Melolontha melolontha* or an area with no or only low density of *M. melolontha* over the last decades (Huber, Bont, et al., 2016). The larvae of *M. melolontha* are one of the major native root herbivores of *T. officinale* and can severely impair the plant (Hauss & Schütte, 1976; Huber, Bont, et al., 2016). Due to the low mobility of larvae and habitat fidelity of adults, local populations of *M. melolontha* remain stable over long periods of time, and in a previous study we found that past *M. melolontha* abundance is a good predictor of present *M. melolontha* pressure (Huber, Bont, et al., 2016).

### Generation of F2 plants

To investigate heritable variation in latex secondary metabolites, we grew plants from field-collected seeds in a greenhouse for two generations. F2 plants were obtained by controlled hand-pollination between F1 plants of the same population. Initially, we collected one seed head from each of the 15 selected F0 mother plants per field site during the field visits. In August 2016, F1 seeds were germinated in a greenhouse (22 °C day / 18 °C night; 16 h light / 8 h darkness) by sowing 10-15 seeds per collected seed head into a small pot (5.5 cm diameter) filled with moist seedling substrate. After three weeks, three seedlings of each mother plant were transplanted together as a seed family into one bigger pot (13 x 13 x 13 cm) filled with potting soil (5 parts field soil, 4 parts peat, and 1 part sand). As not all seeds germinated, we obtained between 6 and 15 seed families per population, with a total of 931 pots. One week after transplantation, pots were put outside for growth and natural vernalisation during fall of 2016. In November 2016, pots were transported into a semi-controlled greenhouse (temperature between 15 and 25 °C, 16 h light and 8 h darkness), where flower production started after 1-2 months. Across all populations at least one plant per population produced flowers, however, number of flowering plants varied between populations. To avoid cross-pollination between populations, flower buds were covered with a tea bag before flower opening. Whenever flowers of two plants from different seed families within the same population were open concurrently, tea bags of those flowers were removed, flowers were carefully rubbed against each other for pollination and then covered again with tea bags until seed collection. As triploid plants reproduce clonally, pollination is only necessary for the obligate outcrossing diploids. However, at the time of pollination, ploidy level of F1 plants was unknown, thus, hand-pollination was done for all plants. After collection, seeds were stored at 4°C.

To grow F2 plants for latex analysis, we included all populations for which we could collect seeds from 3-6 different F1 plants per population. We excluded eight of the 63 populations for having fewer than three seed-producing F1 plants. Ploidy level analysis of F1 plants (see below) revealed that populations with mixtures of diploid and triploid cytotypes were frequent (Fig. 1). F1 parent plants were selected according to their ploidy level to represent the ratio of diploid and triploid plants of each population. Rarely occurring tetraploid plants were excluded. For populations with more than 75% diploid plants, all parent plants for the F2 seeds were chosen to be diploid. For populations with 50-75% diploid plants, 2/3 diploid and 1/3 triploid plants were chosen as parent plants. For populations with 25-50% diploid plants, 1/3 diploid and 2/3 triploid plants were chosen as parent plants. For population with less than 25% diploid plants, all parent plants were chosen to be triploid. Seeds were germinated in a greenhouse (22 °C day / 18 °C night; 16 h light / 8 h darkness) on moist seedling substrate. After three weeks, one seedling per parent was transplanted into a pot (9 cm x 9 cm x 9 cm) filled with garden soil (Selmaterra, Eric Schweizer AG, Thun, Switzerland), resulting in a total of 256 pots. Pots were kept in the greenhouse in a randomized fashion until latex collection.

### Ploidy level

To estimate the percentage of diploid, triploid and tetraploid plants per population, DNA ploidy level of F1 plants was determined by flow cytometry. A CyFlow Cube (Sysmex Partec GmbH, Goerlitz, Germany) with a Partec CyStain UV precise P kit (ref. 05-5002) was used following the manufacturer’s instructions. Ploidy level was determined using fresh tissue from one leaf per plant per pot. As external standard, leaf tissue of *T. officinale* plants with known ploidy level (diploid or triploid) was used. Ploidy levels of F1 plants were estimated by comparing the sample peak to the standard peak. Approximately 1500 nuclei were measured per sample.

### Latex collection and chemical analysis

Latex from F0 plants was collected in each field site from the 15 plants that were selected for seed collection. Latex from F2 plants was collected in the greenhouse when plants where three months old. To obtain taproot latex, each plant was cut 0.5 cm below the tiller and 2 μl of the exuding latex was pipetted immediately into 200 μl methanol. After returning to the lab, latex samples were stored at -20 °C until processing. For extraction of latex secondary metabolites, tubes were vortexed for 10 min, ultrasonicated for 10 min, centrifuged at 4 °C and 14000 rpm for 20 min and supernatants were stored at −20 °C. For F0 plants, pooled samples per population were made for chemical analysis by mixing 10 μl of methanol extract from each of the 15 plants per population in an Eppendorf tube. For F2 plants, methanol extracts from individual samples were used for chemical analysis and average concentrations per population were calculated afterwards. Relative concentrations of TA-G, di-PIEs and tri-PIEs were determined as described in Bont et al. (2017). In brief, methanol extracts were injected into an Acquity UPLC-PDA-MS (Waters, Milford MA, USA) with electrospray ionization in positive mode, consisting of an ultra-performance liquid chromatograph (UPLC) coupled to a photodiode array detector (PDA) and a single quadrupole mass detector (QDa). For quantification, peak areas were integrated at 245 nm for TA-G and at 275 nm for di- and tri-PIES, while concurrently recorded characteristic mass features were used to confirm compound identities.

### Statistical analysis

To disentangle heritable variation in latex secondary metabolites and variation due to environmental plasticity, we performed linear regression analyses within each class of secondary metabolites to investigate how the chemistry of F2 plants is related to the chemistry of F0 plants. If, within a class of secondary metabolites, the linear regression between F2 plants and F0 plants was statistically significant, we used the slope of this regression as an approximate estimate for narrow-sense heritability (pseudo-*h*^2^) (Falconer, 1981). We also compared patterns of covariation among latex metabolites by testing for correlations between the three metabolite classes within F0 plants and within F2 plants separately.

To investigate the effects of climate, soil and belowground herbivores on the latex profile of *T. officinale,* we applied a set of linear mixed-effects models across all populations. As the climate dataset showed multicollinearity, we first applied a principal component analysis (PCA) on the ten selected meteorological variables to reduce dimensionality. We scaled the variables and used the function ‘prcomp’ in R to extract the values of the first (climPCA1) and the second (climPCA2) axis. Both axes together explained 76% of the variation in the original climate data and were then used to represent the climatic conditions of each population. In order to analyse whether the climatic conditions are linked with the geographical localisation of the populations, we tested for correlations between the climatic PCA axes and altitude, latitude and longitude of the populations. Further, we tested for correlations among the variables that represent climatic conditions, soil geochemical properties and root herbivore abundance. For linear mixed-effects model analysis, we used the function ‘lmer’ from the package ‘lme4’ (Bates, Maechler, Bolker, & Walker, 2015) to fit models for each *T. officinale* generation (F0 and F2) and each latex secondary metabolite class (TA-G, di-PIEs, tri-PIEs) separately. The response variable was the population mean of the respective secondary metabolite class and climPCA1, climPCA2, humus content, soil pH, *M. melolontha* area (yes / no) and number of belowground herbivores per plant of each population were used as fixed factors. The categorical percentages of diploid plants per population (0-24%, 25-74%, 75-100%) were added as a random factor, as sexual diploids and asexual triploids have a different reproduction system, which can affect the heritability of a trait. Interaction terms were not included in the models. If necessary, response variables were log-transformed to improve distribution of variance. Effect sizes were estimated using restricted maximum likelihood (REML). Models were validated using ‘plotresid’ from the package ‘RVAideMemoire’ (Hervé, 2018). The significances of the fixed effects were estimated by Wald chi-square tests using the function ‘Anova’ from the package ‘car’ (Fox & Weisberg, 2011). In order to facilitate the visualization of significant model effects, we additionally performed separate linear regressions for each significant fixed factor in the lmer models. Similar results were obtained with models for values of individual plants, which included population as additional random factor, compared to models for mean values per population.

A map of all populations was created with ‘ggmap’ (Kahle & Wickham, 2013), ‘viridis’ (Garnier, 2018) and ‘ggsn’ (Baquero, 2017). Results were visualized using ggplot2 (Wickham, 2016) and factoextra (Kassambara & Mundt, 2017). All statistical analyses were performed in R 3.4.0 (R Core Team, 2017).

## Results

### Latex metabolite production varies greatly across populations

Both in natural habitats (F0 plants) and under greenhouse conditions (F2 plants), the production of latex secondary metabolites varied greatly across *T. officinale* populations (Figs. S1-S3). In the field, TA-G differed almost 19-fold (Fig. S1a), di-PIEs more than sevenfold (Fig. S2a) and tri-PIEs more than 200-fold (Fig. S2c) among populations. Under greenhouse conditions, mean concentrations of latex secondary metabolites were increased but varied less among populations. Nonetheless, average concentrations of TA-G still differed almost threefold (Fig. S1b; average calculated without one population that produced almost no TA-G), di-PIEs more than threefold (Fig. S2b) and tri-PIEs more than 150-fold (Fig. S3b) among populations. The mean production of latex metabolites by F2 plants was partially predicted by mean concentrations in F0 plants for TA-G (*R*^2^ = 0.12, *F*_(1,52)_ = 6.07, *P* = 0.01, Fig 2a) and tri-PIEs (*R*^2^ = 0.45, *F*_(1,52)_ = 41.78, *P* < 0.001, Fig. 2c). This suggests that heritable genetic variation contributed to the variation in these secondary metabolites, with estimated narrow-sense heritabilities (slope ± SE) of 0.39 ± 0.15 for TA-G and 0.77 ± 0.12 for tri-PIEs. For di-PIEs, no statistically significant influence of the production of plants growing in natural habitats (F0) on the production of plants growing under greenhouse conditions (F2) was found (Fig. 2b).

**Figure 2.**
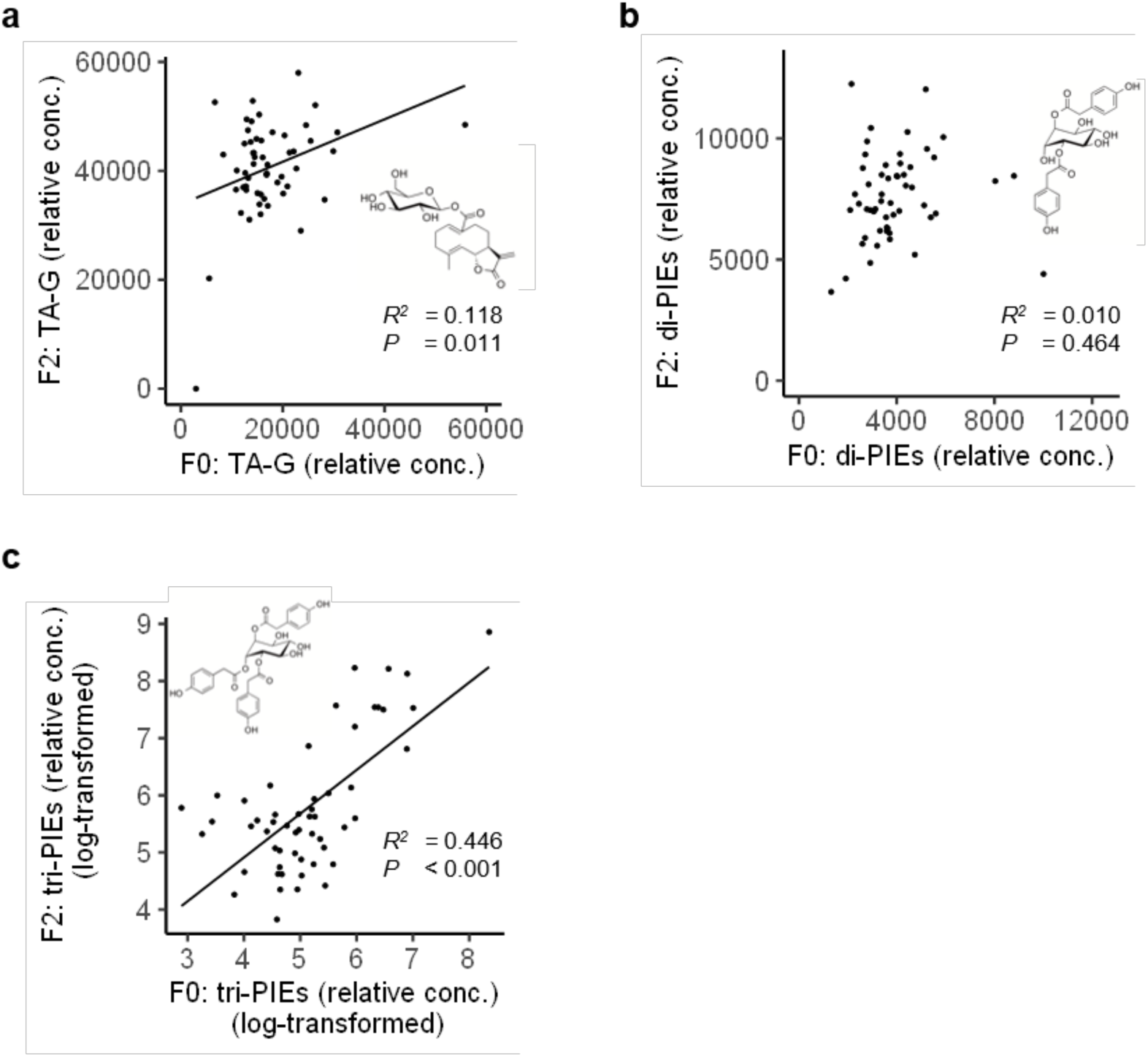
Relationship between the latex chemistry of the plants growing in natural habitat (F0) and the plants growing in greenhouse conditions (F2). Regression analyses were performed separately for TA-G (a), di-PIEs (b) and tri-PIEs (c). Each dot represents one of the 63 sampled *T. officinale* populations. *R*^2^- and *P-*values are displayed and, for statistically significant regressions (*P* < 0.05), linear regression lines are shown. TA-G: taraxinic acid ß-D-glucopyranosyl ester; di-PIEs: di-4-hydroxyphenylacetate inositol esters; tri-PIEs: tri-4-hydroxyphenylacetate inositol esters.

### Environmental plasticity contributes to variation in latex secondary metabolites

In order to examine covariation of latex metabolite concentrations and to assess the additional component of environmental plasticity besides heritable variation in the latex profiles, we tested for pairwise correlations among the three classes of secondary metabolites for plants growing in natural habitat (F0) and plants growing in the greenhouse (F2). The analysis revealed that for F0 plants, TA-G, di-PIEs and tri-PIEs were significantly positively correlated with each other (TA-G and di-PIEs: Pearson’s *r* = 0.76, *P* < 0.001; TA-G and tri-PIEs: Pearson’s *r* = 0.66, *P* < 0.001; di-PIEs and tri-PIEs: Pearson’s *r* = 0.42, *P* < 0.001; Fig 3a). In contrast, only TA-G and di-PIEs were positively correlated for F2 plants (Pearson’s *r* = 0.38, *P* = 0.004, Fig 3b), while TA-G and tri-PIEs were not significantly correlated (Pearson’s *r* = 0.04, *P* = 0.78, Fig 3b) and di-PIEs and tri-PIEs were negatively correlated (Pearson’s *r* = -0.41, *P* = 0.002, Fig 3b). This finding indicates that the regulation of latex secondary metabolites is plastic in response to the environment and not solely determined by genetic, heritable variation.

**Figure 3.**
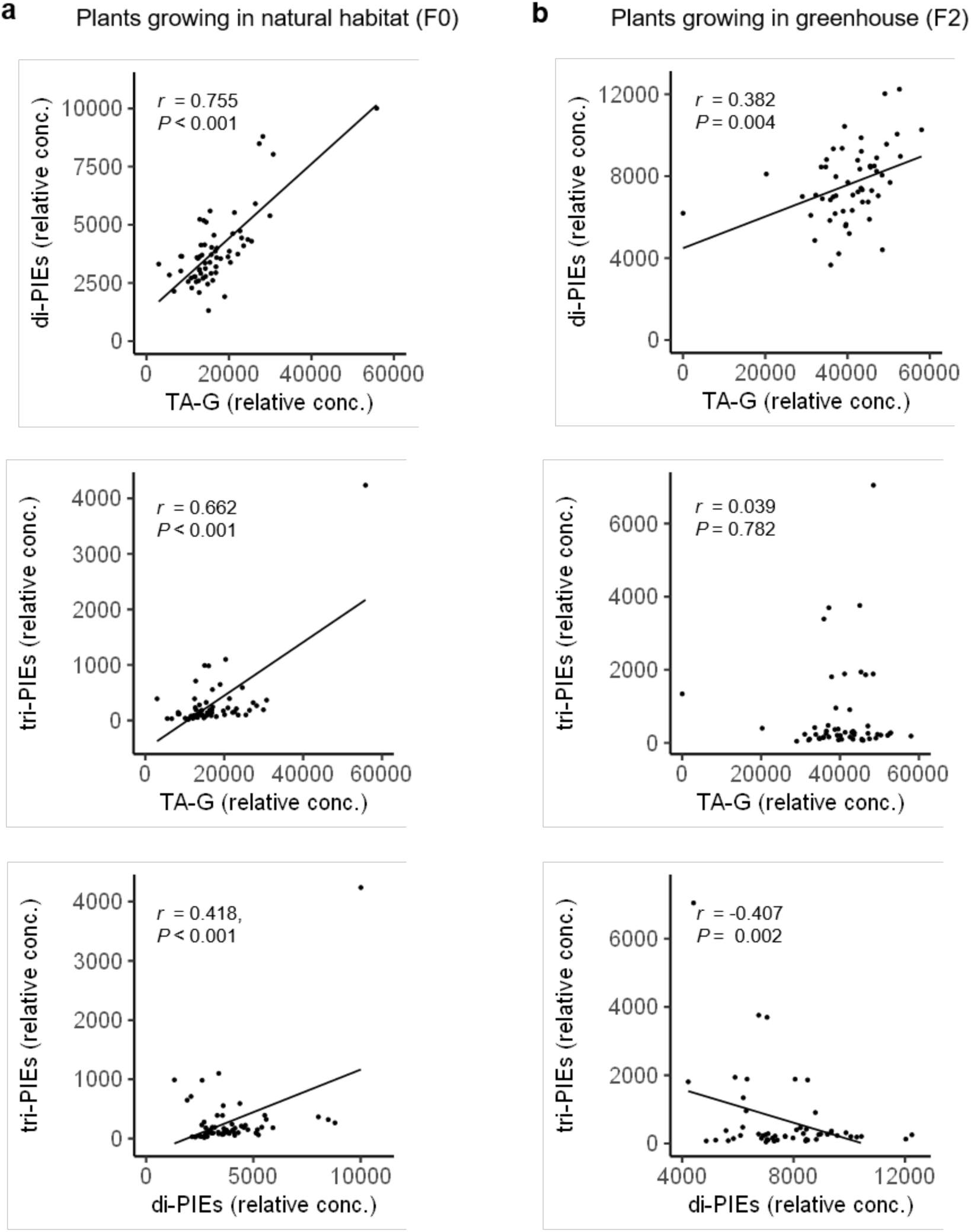
Covariation of the three analysed classes of latex secondary metabolites for plants growing in natural habitat (a) and plants growing in greenhouse (b). Each dot represents one of the 63 sampled *T. officinale* populations. Pearson’s *r* and corresponding *P-*values are displayed. For statistically significant correlations (*P* < 0.05), estimated linear relationships are visualized with solid lines. TA-G: taraxinic acid ß-D-glucopyranosyl ester; di-PIEs: di-4-hydroxyphenylacetate inositol esters; tri-PIEs: tri-4-hydroxyphenylacetate inositol esters.

### Principal components of climatic niches correlate with altitude and latitude

To explore the effect of abiotic and biotic factors on the latex profile of *T. officinale*, we characterized the environmental conditions of the 63 populations using climate, soil and herbivory variables and tested for their impact on latex secondary metabolites. For soil and herbivory measures we included two variables each, whereas the climatic data consisted of ten in parts strongly correlated meteorological variables (Table 1) and was subjected to dimensionality reduction by PCA, resulting in two climatic components (climPCA1, climPCA2). climPCA1 explained 52.6% of the total variation in climatic variables, and climPCA2 explained a further 23.4% of the total variation (Fig. 4a). Loading scores indicate that climPCA1 was primarily determined by variation in annual temperature (Table 1, Fig. 4a), while climPCA2 was determined by sunshine intensity and inversely correlated with precipitation (Table 1, Fig. 4a). The correlation analysis with geographical variables revealed that climPCA1 was highly correlated with altitude (Pearson’s *r* = -0.95, *P* ≤ 0.001, Fig. 4b), whereas climPCA2 was correlated with latitude (Pearson’s *r* = -0.53, *P* ≤ 0.001, Fig. 4b). Neither climPCA1 nor climPCA2 were correlated with longitude (Pearson’s *r* = 0.01, *P* = 0.96 respectively Pearson’s *r* = -0.14, *P* = 0.28). No correlations were found among the climate, soil and herbivory parameters that were selected to represent the environmental conditions of the populations (Table S2).

**Figure 4.**
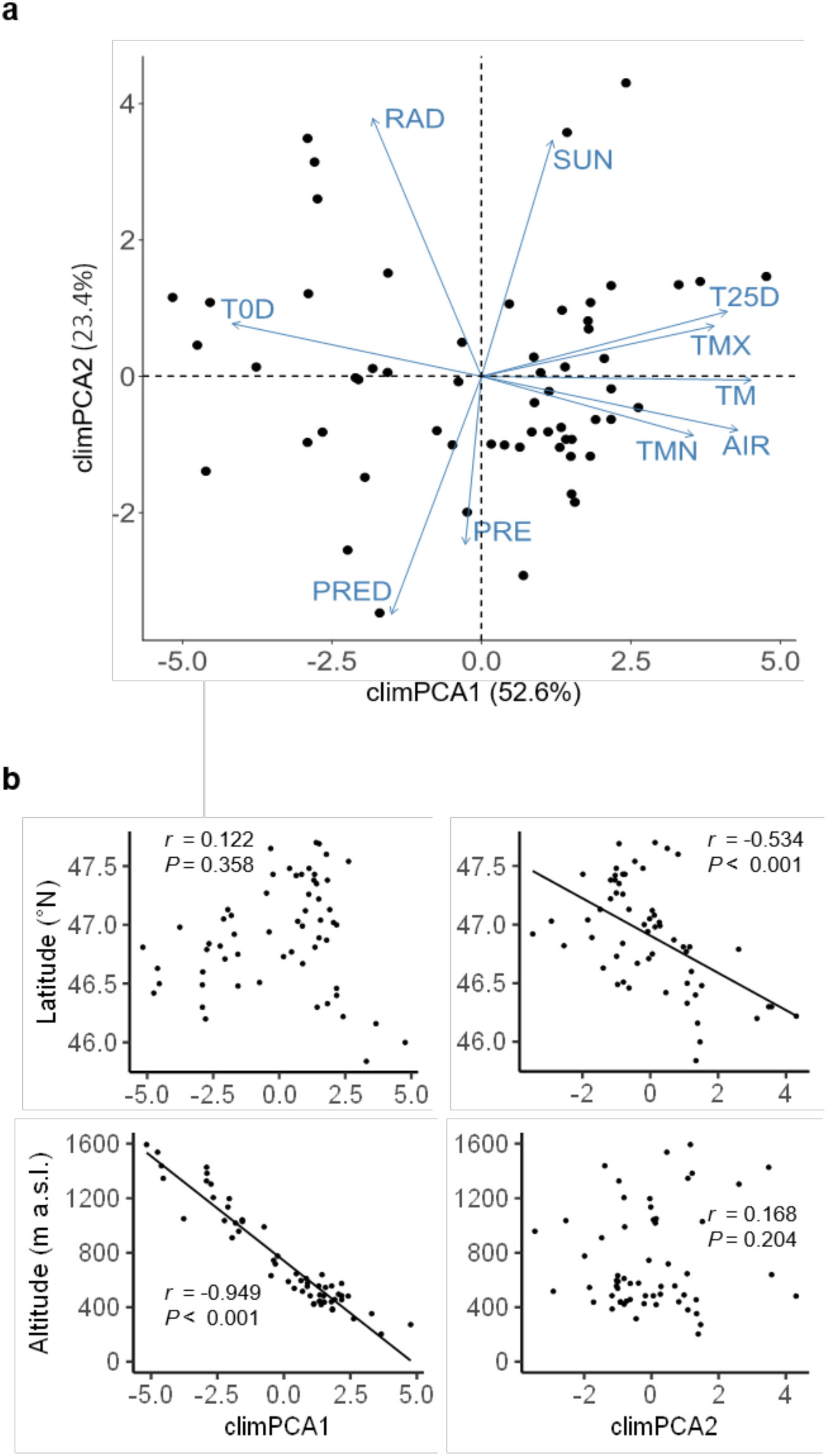
Climatic conditions of the sampled *T. officinale* populations. (a) Biplot of a principal component analysis of the climatic conditions of the populations. The first two axes (climPCA1, climPCA2) are shown, which explain 76% of cumulative variance. Each dot represents one of the 63 sampled populations. Blue vectors represent the climatic variables. AIR: annual mean of air pressure; PRED: annual precipitation; RAD: annual mean of global radiation; SUN: annual sunshine duration; TM: annual mean temperature; TMN: annual mean of the minimal temperature per day; TMX: annual mean of the maximal temperature per day; T0D: days per year with a minimal temperature below 0 °C; T25D: Days per year with maximal temperature over 25 °C. (b) Pearson correlations of the first and second principal components (climPCA1, climPCA2) of the climatic conditions with the position parameters (altitude and latitude) of the sampled *T. officinale* populations. Each dot represents one of the 63 sampled populations. Linear correlation lines and corresponding *r*- and *P-*values of correlations are shown.

### Heritable variation of latex profile is associated with climate

Using a set of linear mixed-effects models, we tested for shaping effects of climatic, soil and herbivory parameters on the concentrations of latex secondary metabolites across all populations in both plants growing in natural habitat (F0) and plants growing in the greenhouse (F2). We found similar patterns for both generations of plants, which confirms a high degree of heritable variation in latex profiles (Table 2, Fig. 5). The mixed model analyses further revealed distinct effects of the environmental variables depending on the class of secondary metabolite (Table 2). The concentration of TA-G was negatively associated with climPCA2 alone (LMEM, *χ*^2^ = 8.98, *P* = 0.003, Fig. 5a for F0 and *χ*^2^ = 10.03, *P* = 0.001, Fig. 5b for F2, Table 2). As climPCA2 was positively correlated with annual sun intensity and negatively correlated with latitude (Table 1, Fig. 4b), this suggests that populations from the North of Switzerland produced more TA-G than populations from the sun-intense regions in the South of Switzerland (Fig. 5). Interestingly, di-PIEs were not affected by climPCA2, but negatively associated with climPCA1 (LMEM, *χ*^2^ = 6.10, *P* = 0.014, Fig. 5a for F0 and *χ*^2^ = 0.11, *P* = 0.001, Fig. 5b for F2, Table 2). climPCA1 was positively correlated with annual temperature and negatively correlated with altitude (Table 1, Fig. 4b), thus, plants from populations growing at high altitude in colder areas produced more di-PIEs than plants from populations growing in warmer regions at lower altitude (Fig. 5). Mixed model analysis further revealed that neither soil parameters nor belowground herbivore abundance were significantly linked to TA-G or di-PIE production. The number of belowground herbivores per plant was weakly negatively associated with the concentration of tri-PIEs for plants from natural habitats (F0) (LMEM, *χ*^2^ = 3.91, *P* = 0.048, Table 2, Fig. 5a). However, a scatter plot revealed that this result was largely driven by two outlier populations (Fig. 5a).

**Figure 5.**
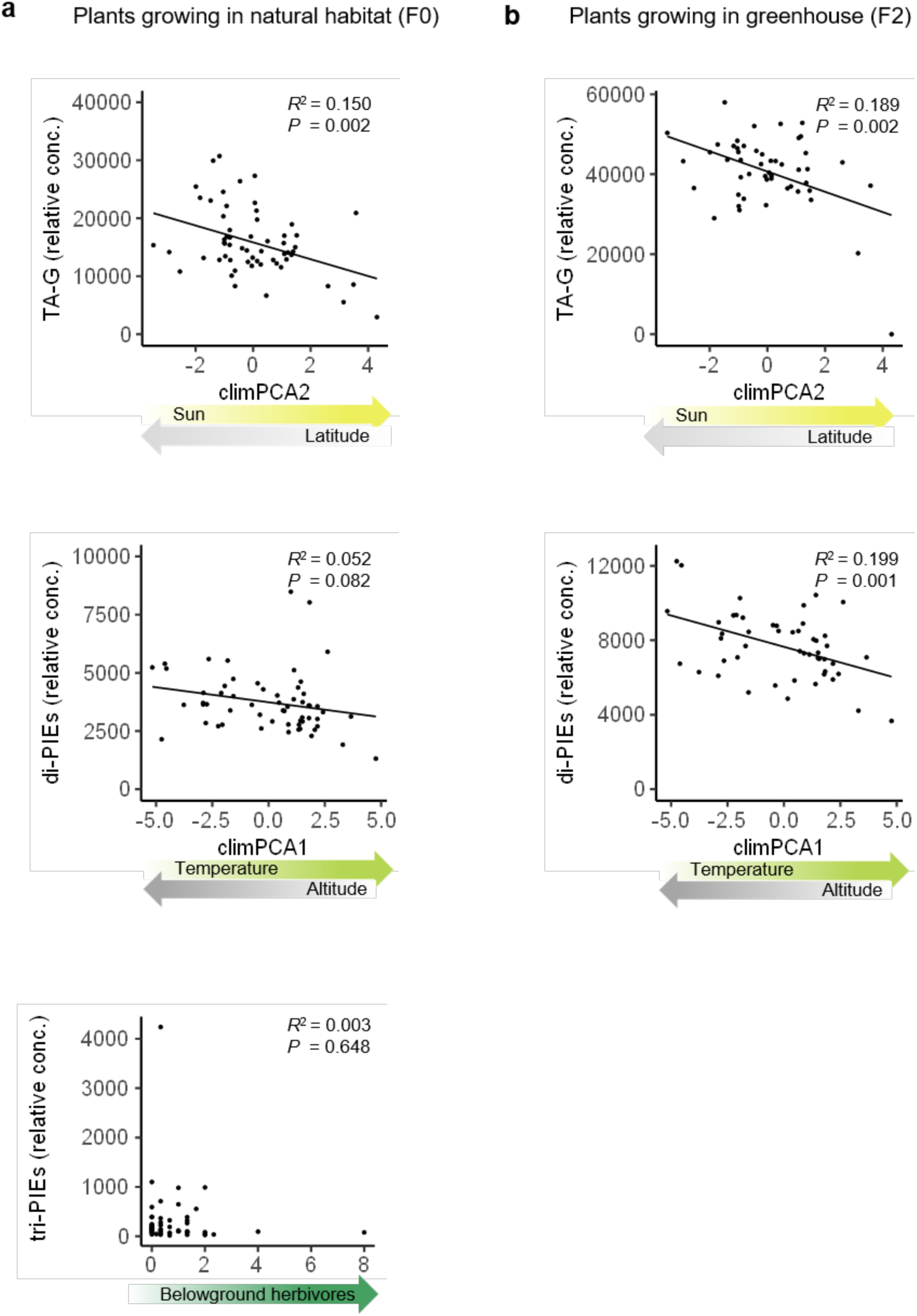
Visualization of relationships between the significant effects of the mixed-effects model and the corresponding class of latex secondary metabolites for plants growing in natural habitat (a) and plants growing in greenhouse (b). Each dot represents one of the 63 sampled *T. officinale* populations. Linear regression lines and *R*^2^- and *P-*values of the regressions are shown. Coloured arrows illustrate intensity gradients of the corresponding environmental factors. Note that these linear regressions are different from the mixed-effects models. The regressions in this figure simplify the relationships between latex metabolites and environmental factors for illustrative purposes, while the mixed-effect models in Table 2 examine the associations in a more powerful modelling framework. TA-G: taraxinic acid ß-D-glucopyranosyl ester; di-PIEs: di-4-hydroxyphenylacetate inositol esters; tri-PIEs: tri-4-hydroxyphenylacetate inositol esters.

**Table 2.**
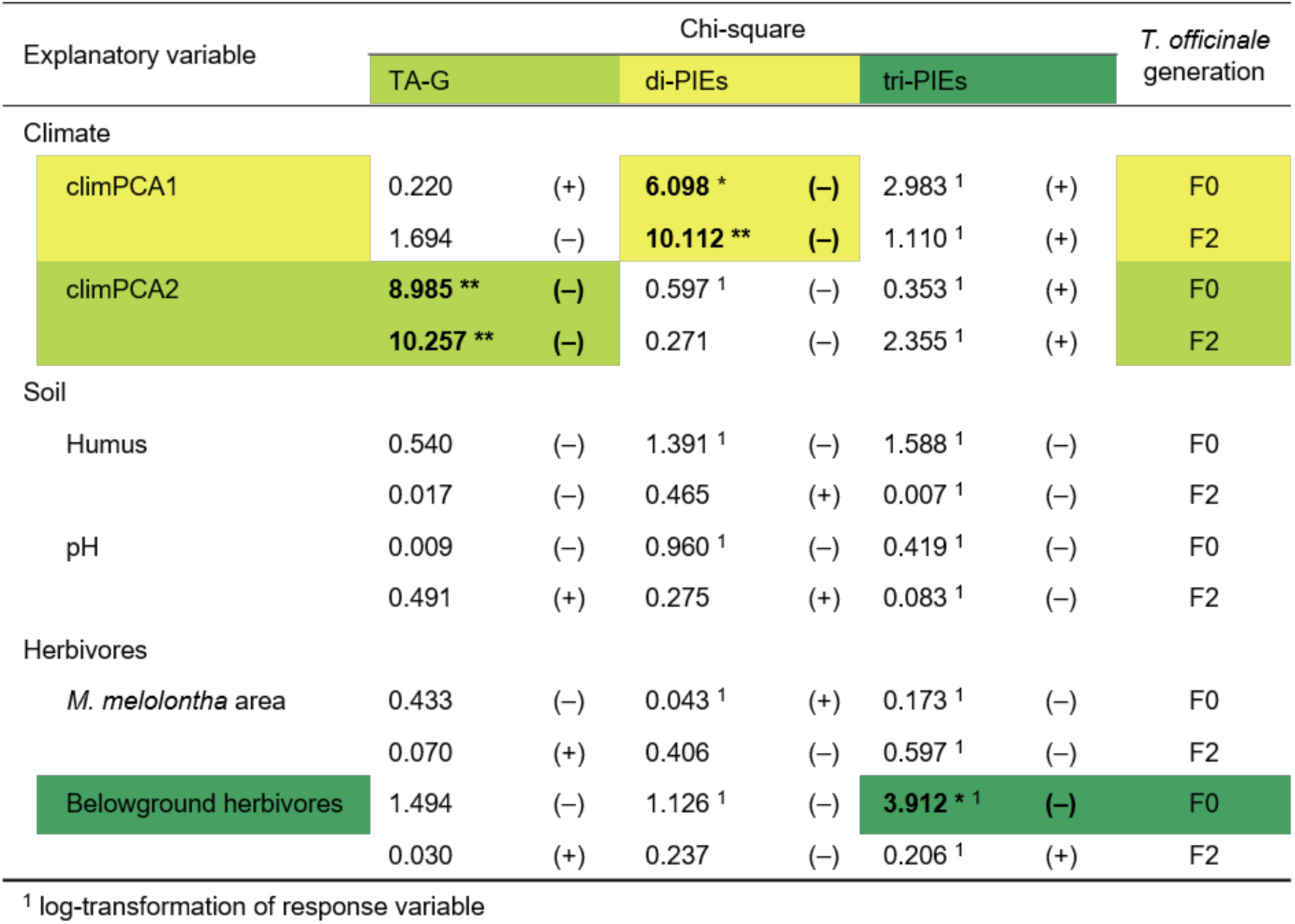
Summary of results from mixed-effects models that relate latex secondary metabolite concentrations to variables representing the environmental conditions of the investigated *T. officinale* populations. Models have been established separately for population means of each class of secondary metabolite as response variable and separately for each *T. officinale* generation. Environmental variables (climPCA1, climPCA2, humus, pH, *M. melolontha* area, belowground herbivores) were included as fixed effects and the percentage of diploid plants per population as random effect. Significances of the fixed effects were estimated by Wald chi-square tests. Chi-square values are displayed as numbers, whereas plus and minus signs refer to the effect direction of each factor. Statistically significant effects with *P* < 0.05 (\*\*\**P* < 0.001; \*\**P* < 0.01; \**P* < 0.05) are displayed with bold numbers and highlighted with colour patterns. TA-G: taraxinic acid ß-D-glucopyranosyl ester; di-PIEs: di-4-hydroxyphenylacetate inositol esters; tri-PIEs: tri-4-hydroxyphenylacetate inositol esters.

| Explanatory variable | Chi-square |  |  |  |  |  | *T. officinale*<br>generation |
| --- | --- | --- | --- | --- | --- | --- | --- |
|  | TA-G |  | di-PIEs |  | tri-PIEs |  |  |
| Climate |  |  |  |  |  |  |  |
| climPCA1 | 0.220 | (+) | 6.098 * | (-) | 2.983 <sup>1</sup> | (+) | F0 |
|  | 1.694 | (-) | 10.112 ** | (-) | 1.110 <sup>1</sup> | (+) | F2 |
| climPCA2 | 8.985 ** | (-) | 0.597 <sup>1</sup> | (-) | 0.353 <sup>1</sup> | (+) | F0 |
|  | 10.257 ** | (-) | 0.271 | (-) | 2.355 <sup>1</sup> | (+) | F2 |
| Soil |  |  |  |  |  |  |  |
| Humus | 0.540 | (-) | 1.391 <sup>1</sup> | (-) | 1.588 <sup>1</sup> | (-) | F0 |
|  | 0.017 | (-) | 0.465 | (+) | 0.007 <sup>1</sup> | (-) | F2 |
| pH | 0.009 | (-) | 0.960 <sup>1</sup> | (-) | 0.419 <sup>1</sup> | (-) | F0 |
|  | 0.491 | (+) | 0.275 | (+) | 0.083 <sup>1</sup> | (-) | F2 |
| Herbivores |  |  |  |  |  |  |  |
| *M. melolontha* area | 0.433 | (-) | 0.043 <sup>1</sup> | (+) | 0.173 <sup>1</sup> | (-) | F0 |
|  | 0.070 | (+) | 0.406 | (-) | 0.597 <sup>1</sup> | (-) | F2 |
| Belowground herbivores | 1.494 | (-) | 1.126 <sup>1</sup> | (-) | 3.912 * <sup>1</sup> | (-) | F0 |
|  | 0.030 | (+) | 0.237 | (-) | 0.206 <sup>1</sup> | (+) | F2 |

<sup>1</sup> log-transformation of response variable

## Discussion

Plant secondary metabolites can vary considerably between populations, but the contribution of the environment in shaping secondary metabolite profiles of plants by driving selection or environmental plasticity is poorly understood. Our work shows that heritable variation in latex secondary metabolites of 63 *T. officinale* populations across Switzerland is strongly linked to climatic conditions, but not to soil properties or belowground herbivore abundance. Here, we discuss the implications of these findings from ecological and physiological points of view.

The *T. officinale* populations included in this study were evenly distributed across Switzerland and spanned an elevation gradient from 200 – 1600 m a.s.l. For Switzerland, the occurrence of both sexual diploid and asexual triploid dandelions has been reported, with diploids being found at elevations higher than 700 m a.s.l and triploids being found in lower areas (Calame & Felber, 2000). However, our ploidy analysis could not confirm such an elevation threshold, as we found diploids and triploids at all elevations and frequently observed populations with mixed cytotypes. Calame & Felber (2000) analysed the distribution of *T. officinale* cytotypes along two elevation gradients of different regions (Jura and Alps), whereas in our study we did not sample directly along elevation gradients, but analysed a broader spectrum of populations from all over Switzerland. The differences in the reported cytotype distributions may thus be due to the different sampling ranges.

As a chemical interface between plants and their environment, some plant secondary metabolites vary highly in concentration and composition with changing abiotic conditions (Holopainen et al., 2018; Jakobsen & Olsen, 1994; Selmar & Kleinwächter, 2013). Fluctuating patterns of sesquiterpene lactones and phenolics produced by *Tithonia diversifolia,* for instance, correlate with seasonal changes in temperature and rainfall (Sampaio, Edrada-Ebel, & Da Costa, 2016). Our results show that the concentrations of both TA-G and di-PIEs in the latex of *T. officinale* are strongly associated with the climatic conditions of the population origins, and thus emphasize the central role of abiotic conditions for shaping latex composition. Interestingly, despite the structural similarity of di-PIEs and tri-PIEs, we found no effect of climate on tri-PIEs, which points towards distinct functions and regulations of the two groups of secondary metabolites. Climatic effects on latex profiles were consistent for plants growing in natural habitat and for plants growing under greenhouse conditions two generations later, although altered correlations among compounds in field- or greenhouse-grown plants suggest some degree of environmental plasticity in latex secondary metabolites. Nonetheless, this indicates that variation in latex profiles is at least in part heritable and suggests that climatic conditions exert direct or indirect selection pressure on latex secondary metabolites in *T. officinale*.

Variation in plant defenses between and within species are often hypothesized to follow geographical gradients, which in turn are speculated to be linked to herbivore pressure (Anstett et al., 2018; Moles et al., 2011; Pratt et al., 2014). Both TA-G and di-PIEs have been shown to be involved in herbivore defense (Agrawal et al., 2018; Bont et al., 2017; Huber, Bont, et al., 2016; Huber, Epping, et al., 2016). Our results confirm a link of both metabolites with geographical gradients, as the climatic variables, which influence TA-G and di-PIEs, are strongly linked to altitude and latitude. We found more di-PIEs in plants from high altitudes with lower temperatures, and more TA-G in plants from areas with less sun in the North of Switzerland. However, as herbivore pressure varied independently of climatic variables, we found no evidence for a major role of belowground herbivores in the observed genetically based variation in latex secondary metabolites, contrary to our expectations. Nevertheless, we cannot rule out potential hidden effects of herbivores on latex metabolites. The climatic variables used in our study include 20 years of weather data at very high resolution, whereas our herbivore variables either captured a single snapshot in time, or relied on rough, potentially inaccurate historic estimates. Thus, the two herbivore variables may have failed to accurately represent the actual herbivore abundances of the past (Huber, Bont, et al., 2016). Furthermore, herbivore effects could manifest themselves through interactions with climate conditions or other variables. Plant responses to abiotic and biotic stresses are controlled by the same interactive hormonal network, hence combined stresses may lead to complex hormonal interactions (Nguyen, Rieu, Mariani, & van Dam, 2016). In maize, for example, root herbivory induces hydraulic changes in the leaves and triggers abscisic acid (ABA) accumulation (Erb et al., 2011). ABA again is essentially involved in regulating responses to abiotic stresses and stimulates for instance stomata closure, which is crucial for limiting desiccation (Daszkowska-Golec & Szarejko, 2013). Thus, by influencing the water balance of the plant, root herbivores may indirectly affect the plant’s response to abiotic conditions. However, we could not test for interactive effects of the environmental variables included in our study, as the full set of required tests would have exceeded the appropriate number of model parameters compared to the number of included populations in our analysis. Thus, to shed light on this topic, further experiments are needed, which test specifically the interacting effect of climatic conditions and herbivore pressure on secondary metabolites at large-scale environmental variation. However, given the different and specific associations between PIEs, TA-G and climatic conditions and the inverse relationship between climate parameters, expected herbivore attack rates and secondary metabolite concentrations, we consider it unlikely that the climate effects observed here are the indirect result of climate-mediated herbivory alone.

There is growing evidence that many secondary metabolites are highly multifunctional and serve defensive, ecological and physiological functions at the same time, which minimizes the plant’s fitness costs for biosynthesis and maintenance of metabolites (Bednarek & Osbourn, 2009; Neilson et al., 2013). Maize plants, for instance, use the same benzoxazinoid secondary metabolites for iron uptake, protection against generalist herbivores, and defence signalling (Hu et al., 2018; B. Li et al., 2018; Maag et al., 2016). For another important class of defensive secondary metabolites, glucosinolates, it has been shown that the metabolite 3-hydroxypropylglucosinolate has signalling capacity and inhibits root growth and development by the evolutionary old TOR (Target of Rapamycin) pathway (Malinovsky et al., 2017). In addition, recent work shows that aliphatic glucosinolates have an important role in drought conditions by regulating stomatal aperture, thus providing evidence that glucosinolates are also involved in abiotic stress tolerance (Salehin et al., 2019). Our finding that TA-G and di-PIEs are strongly associated to climatic conditions suggests potential alternative functions of those latex secondary metabolites, and latex itself, in addition to the previous reported roles in herbivore defence (Agrawal et al., 2018; Huber, Bont, et al., 2016). Both TA-G and di-PIE concentrations were correlated with each other, but each compound class was affected by distinct climate variables: TA-G was mainly associated with sun and rain intensity (climPCA2), whereas di-PIEs were mainly associated with temperature (climPCA1). Hence, TA-G may be involved in moisture regulation or linked to physiological processes that are associated with light availability, whereas di-PIEs may play a role in temperature-sensitive processes. Of course these are highly speculative arguments based on correlational data, and manipulative experiments are needed to further explore the role of latex secondary metabolites in climate-mediated plant physiology.

Studies of inter- and intraspecific plant trait variation across environmental gradients, such as those related to latitude and elevation, have been receiving increasing attention (Anstett et al., 2018; Hahn et al., 2018; Pellissier, Roger, Bilat, & Rasmann, 2014; Woods et al., 2012). Although such studies are beyond doubt important and useful to test classic theories predicting herbivore defence (Anstett et al., 2015; Moles et al., 2011) and resource allocation patterns (Helsen et al., 2017; Kooyers, Greenlee, Colicchio, Oh, & Blackman, 2015), they also have to cope with the difficulty of potentially hidden dynamics along gradients. Changes in abiotic and biotic factors may be correlated and interconnected to changes in geographical location, which complicates the disentangling of environmental impacts on plant traits (Hahn et al., 2018; Johnson & Rasmann, 2011). Our results emphasize the importance of considering multiple environmental factors when studying biogeographical patterns of plant traits, and of sampling a large set of natural genotypes across a wide range of environments. We propose to include the possibility of multifunctionality of secondary metabolites into the framework of studies that explore trait variation in plant defences, as patterns of defence variation may be explained by alternative additional functions of plant secondary metabolites.

## Acknowledgements

We thank Armin Komposch, Cyrill Delfgou, Gabriel Ulrich and Zephyr Züst for their help during fieldwork. We are grateful to the gardeners of the University of Bern as well as to Valentin Pulver, Tala Bürki, Andrea Bonini, Robin Bautzmann, Conradin Lutz, Gabriel Zala and Yves Garnier for their help with experiments. We thank Pierre Mateo for drawing chemical structures. Meteorological data was provided by MeteoSwiss, the Swiss Federal Office for Meteorology and Climatology. This study was supported by the Swiss National Science Foundation (Grant No. 153517) and the Seventh Framework Programme for Research and Technological Development of the European Union (FP7 MC-CIG 629134).

## Authors’ contributions

ZB, TZ, MH and ME designed the study. ZB collected the data. ZB, TZ and ME analysed and interpreted the data. ZB and ME wrote the first draft of the manuscript. All authors contributed to the final version of the manuscript.

## Data accessibility

All data supporting this study will be stored in the Dryad Digital Repository and the data DOI will be included in the manuscript.

