## Supplementary Material for "Variation in root secondary metabolites is shaped by past climatic conditions"

**Table S1** Descriptive parameters of the 63 sampled *T. officinale* populations are shown including population number, abbreviation of the corresponding weather station, exact position (latitude, longitude, altitude), percentage of diploid plants and land use of the field during the last decades.

| Nr. | Weather station | Latitude (°N) | Longitude (°E) | Altitude (m a.s.l.) | Diploids (%) | Land use |
|---|---|---|---|---|---|---|
| 1 | ABO | 46°29′28″ | 7°33′41″ | 1306 | 100 | Extensive cutting and grazing |
| 2 | AIG | 46°19′51″ | 6°55′40″ | 381 | 13.3 | Extensive cutting and grazing |
| 3 | ALT | 46°53′25″ | 8°37′12″ | 437 | 100 | Middle-intensive cutting and grazing |
| 4 | ANT | 46°37′52″ | 8°35′16″ | 1433 | 100 | Extensive cutting and grazing |
| 5 | BAS | 47°32′32″ | 7°34′45″ | 303 | 100 | Intensive cutting and grazing |
| 6 | BER | 46°59′31″ | 7°27′37″ | 553 | 93.3 | Extensive cutting |
| 7 | BIL | 47°06′45″ | 7°15′30″ | 474 | 0 | Grazing |
| 8 | BLA | 46°25′11″ | 7°49′14″ | 1530 | 100 | Extensive cutting |
| 9 | BRL | 46°59′01″ | 6°37′02″ | 1042 | 27.3 | Grazing |
| 10 | BUS | 47°23′25″ | 8°05′05″ | 388 | 87.5 | Intensive grazing |
| 11 | CDF | 47°04′48″ | 6°48′21″ | 1026 | 85.7 | Middle-intensive cutting and grazing |
| 12 | CGI | 46°23′37″ | 6°13′11″ | 441 | 42.9 | Extensive cutting |
| 13 | CHD | 46°28′58″ | 7°08′07″ | 1117 | 100 | Middle-intensive cutting and grazing |
| 14 | CHM | 47°03′10″ | 6°58′49″ | 1132 | 100 | Extensive cutting and grazing |
| 15 | CHU | 46°52′41″ | 9°32′14″ | 593 | 100 | Intensive cutting and grazing |
| 16 | CLA | 46°27′00″ | 6°54′06″ | 528 | 100 | Intensive cutting |
| 17 | COM | 46°26′06″ | 8°57′39″ | 462 | 93.3 | Intensive cutting |
| 18 | DAV | 46°48′40″ | 9°51′01″ | 1568 | 92.9 | Extensive cutting and grazing |
| 19 | DEM | 47°21′30″ | 7°20′13″ | 419 | 100 | Extensive cutting and grazing |
| 20 | DIS | 46°42′24″ | 8°51′13″ | 1196 | 100 | Extensive cutting and grazing |
| 21 | EBK | 47°16′26″ | 9°06′44″ | 623 | 100 | Intensive cutting and grazing |
| 22 | EIN | 47°07′53″ | 8°45′12″ | 908 | 100 | Middle-intensive cutting and grazing |
| 23 | ELM | 46°55′02″ | 9°10′15″ | 986 | 100 | Middle-intensive cutting and grazing |
| 24 | ENG | 46°49′21″ | 8°24′41″ | 1063 | 100 | NA |
| 25 | FAH | 47°25′43″ | 6°56′53″ | 588 | 35.7 | Grazing |
| 26 | FEY | 46°10′51″ | 7°15′56″ | 998 | 100 | Extensive cutting and grazing |
| 27 | FRE | 46°50′26″ | 6°34′42″ | 1200 | 100 | Middle-intensive cutting and grazing |
| 28 | GLA | 47°02′07″ | 9°03′59″ | 495 | 100 | Intensive cutting and grazing |
| 29 | GRA | 46°46′27″ | 7°05′51″ | 627 | 100 | Extensive cutting and grazing |
| 30 | GRC | 46°11′43″ | 7°50′13″ | 1607 | 100 | NA |
| 31 | GUT | 47°36′06″ | 9°16′46″ | 440 | 100 | Intensive cutting and grazing |
| 32 | HAI | 47°39′06″ | 9°01′53″ | 716 | 50 | Grazing |

**Table S1** continued

| Nr. | Weather station | Latitude (°N) | Longitude (°E) | Altitude (m a.s.l.) | Diploids (%) | Land use |
|---|---|---|---|---|---|---|
| 33 | HLL | 47°41′51″ | 8°28′14″ | 419 | 78.6 | Grazing |
| 34 | INT | 46°40′32″ | 7°52′09″ | 575 | 100 | NA |
| 35 | KLO | 47°28′41″ | 8°31′46″ | 435 | 100 | NA |
| 36 | KOP | 47°07′28″ | 7°36′51″ | 480 | 100 | Extensive cutting and grazing |
| 37 | LAG | 46°56′06″ | 7°47′60″ | 687 | 100 | Extensive cutting and grazing |
| 38 | LUG | 46°01′14″ | 8°55′40″ | 289 | 0 | Intensive cutting and grazing |
| 39 | LUZ | 47°02′31″ | 8°17′02″ | 597 | 100 | Grazing |
| 40 | MAG | 46°09′32″ | 8°55′45″ | 202 | 100 | Extensive cutting and grazing |
| 41 | MER | 46°43′39″ | 8°10′35″ | 592 | 100 | Intensive cutting and grazing |
| 42 | MVE | 46°17′55″ | 7°27′38″ | 1418 | 100 | Extensive cutting |
| 43 | NEU | 47°00′22″ | 6°56′37″ | 634 | 20 | Extensive cutting |
| 44 | PAY | 46°48′16″ | 6°54′25″ | 474 | 100 | Extensive cutting and grazing |
| 45 | PIO | 46°30′53″ | 8°41′17″ | 990 | 100 | NA |
| 46 | PLF | 46°44′24″ | 7°15′46″ | 930 | 100 | Intensive cutting and grazing |
| 47 | RAG | 47°01′00″ | 9°30′08″ | 497 | 100 | Grazing |
| 48 | REH | 47°25′49″ | 8°30′22″ | 440 | 81.8 | Extensive cutting and grazing |
| 49 | RHF | 47°32′39″ | 7°47′54″ | 302 | 13.3 | Middle-intensive cutting |
| 50 | RUE | 47°26′06″ | 7°52′45″ | 605 | 92.9 | Middle-intensive cutting and grazing |
| 51 | SBO | 45°50′39″ | 8°56′43″ | 365 | 0 | Extensive cutting |
| 52 | SCU | 46°47′50″ | 10°17′14″ | 1339 | 76.9 | Extensive cutting and grazing |
| 53 | SHA | 47°39′53″ | 8°35′00″ | 434 | 100 | Middle-intensive cutting and grazing |
| 54 | SIO | 46°13′52″ | 7°18′48″ | 510 | 0 | Extensive cutting |
| 55 | SMA | 47°23′24″ | 8°35′55″ | 465 | 73.3 | Grazing |
| 56 | SMM | 46°35′58″ | 10°25′44″ | 1444 | 100 | Extensive cutting |
| 57 | STG | 47°25′31″ | 9°24′11″ | 780 | 100 | Intensive cutting |
| 58 | TAE | 47°28′50″ | 8°55′04″ | 558 | 100 | Intensive cutting |
| 59 | ULR | 46°30′21″ | 8°18′31″ | 1346 | 21.4 | Extensive cutting and grazing |
| 60 | VAD | 47°08′01″ | 9°31′13″ | 455 | 100 | Extensive cutting |
| 51 | VIS | 46°17′58″ | 7°51′31″ | 643 | 93.3 | Intensive cutting and grazing |
| 62 | WAE | 47°12′30″ | 8°40′33″ | 591 | 100 | Middle-intensive cutting and grazing |
| 63 | WYN | 47°15′08″ | 7°46′42″ | 414 | 100 | Extensive cutting |

**Table S2** Correlations among the variables that represent the environmental conditions of the investigated *T. officinale* populations. Pearson correlations are displayed, except for correlations with *M. melolontha* area, for which correlation coefficients of linear regressions are displayed. All correlations are not statistically significant.

| | climPCA1 | climPCA2 | Humus | Soil pH | *M. melolontha* area | Belowground herbivores |
|---|---|---|---|---|---|---|
| climPCA1 | - | 0 | -0.25 | 0.16 | 0.14 | 0.19 |
| climPCA2 | | - | -0.04 | 0.05 | 0.03 | 0.06 |
| Humus | | | - | 0.11 | 0.06 | -0.14 |
| Soil pH | | | | - | 0.18 | -0.11 |
| *M. melolontha* area | | | | | - | 0.01 |
| Belowground herbivores | | | | | | - |

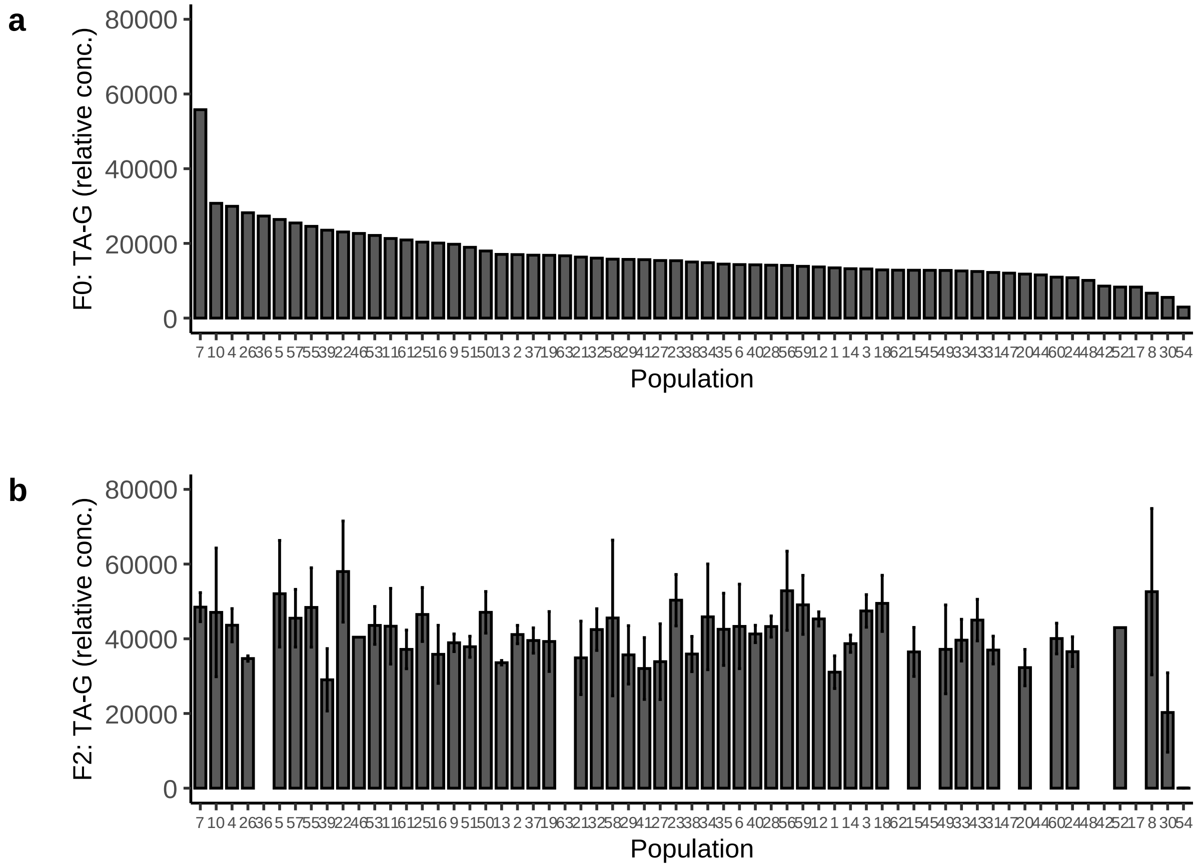


**Figure S1** Concentrations of TA-G in the root latex for each *T. officinale* population. (a) F0 plants growing in natural habitats. Bars represent TA-G concentrations of pooled samples (10 – 15 individuals per population). (b) F2 plants growing in greenhouse conditions. Bars represent average TA-G concentrations of individual plants (*N* = 1-6 per population). Standard errors (± SE) are shown. Note that the population order of (b) corresponds to the population order of (a) and that the values for nine populations growing in greenhouse conditions (b) are not available due to low seed production rate. TA-G: taraxinic acid ß-D-glucopyranosyl ester.

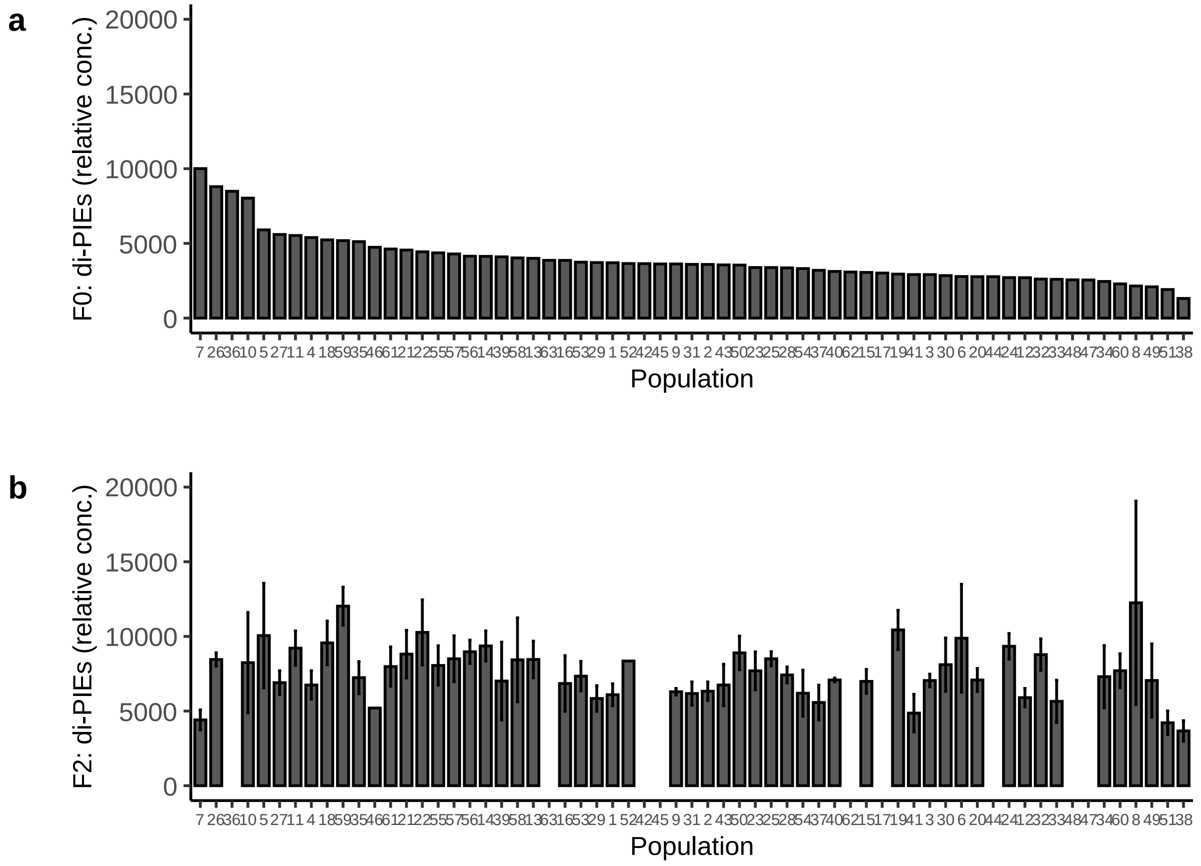

**Figure S2** Concentrations of di-PIEs in the root latex for each *T. officinale* population. (a) F0 plants growing in natural habitats. Bars represent di-PIEs concentrations of pooled samples (10 – 15 individuals per population). (b) F2 plants growing in greenhouse conditions. Bars represent average di-PIEs concentrations of individual plants (*N* = 1-6 per population). Standard errors (± SE) are shown. Note that the population order of (b) corresponds to the population order of (a) and that the values for nine populations growing in greenhouse conditions (b) are not available due to low seed production rate. di-PIEs: di-4-hydroxyphenylacetate inositol esters.

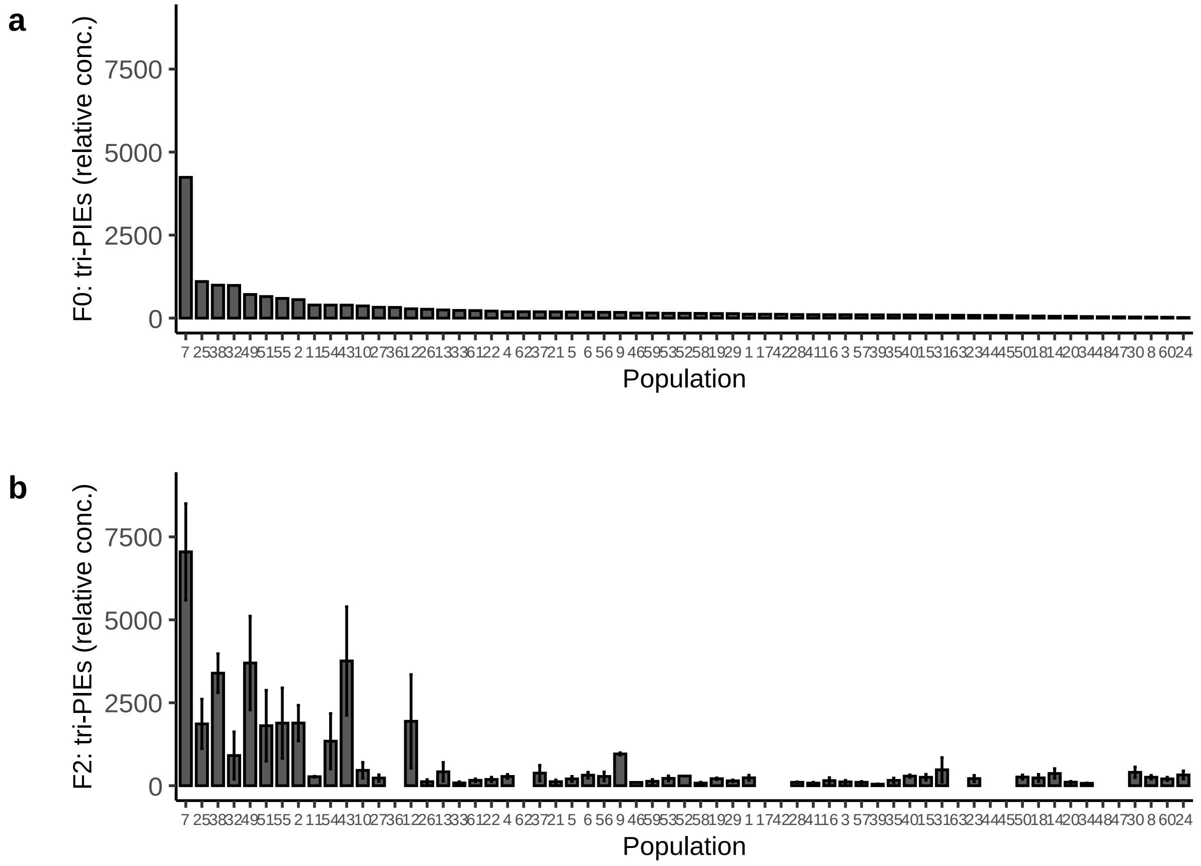


**Figure S3** Concentrations of tri-PIEs in the root latex for each *T. officinale* population. (a) F0 plants growing in natural habitats. Bars represent tri-PIEs concentrations of pooled samples (10 – 15 individuals per population). (b) F2 plants growing in greenhouse conditions. Bars represent average tri-PIEs concentrations of individual plants (*N* = 1-6 per population). Standard errors (± SE) are shown. Note that the population order of (b) corresponds to the population order of (a) and that the values for nine populations growing in greenhouse conditions (b) are not available due to low seed production rate. tri-PIEs: tri-4-hydroxyphenylacetate inositol esters.
